# Marker and readout genes for priming in the Arabidopsis – *Pseudomonas cannabina* pv. alisalensis interaction

**DOI:** 10.1101/2023.07.16.549199

**Authors:** Andrea J. Sistenich, Lisa Fürtauer, Franziska Scheele, Uwe Conrath

## Abstract

After a local infection, the entire plant foliage becomes primed for the superinduction of defense responses after rechallenge. The identity of genes expressed during priming (priming-marker genes) or with hyperexpression in primed plants after rechallenge (priming-readout genes) remained largely unknown. We show in *Arabidopsis thaliana* that genes *AT1G76960* (with unknown function), *AT3G51860* (encoding vacuolar Ca^2+^/H^+^ antiporter CAX3), and *AT3G45860* (cysteine-rich receptor-like protein kinase CRK4) are strongly expressed during *Pseudomonas cannabina* pv. alisalensis-induced priming but not, or much less, in uninfected or challenged plants, or in primed plants after rechallenge. Expression of these genes thus solely marks the primed state for enhanced defense. In contrast to these genes, expression of *AT2G14610* (pathogenesis-related protein PR1), *AT2G32680* (receptor-like protein RLP23), *AT3G25010* (RLP41) and some less expressed loci are activated, about equally, in primed plants before and after rechallenge. They may also serve as marker genes for priming. In further contrast, genes *AT2G39530* (encoding Casparian strip domain-like protein 4D1), *AT2G19190* (flg22-induced receptor-like kinase FRK1), *AT3G28510* (P loop-containing nucleoside triphosphate hydrolases superfamily protein) and some less strongly expressed genes are not, or only faintly activated in uninfected plants, during priming and upon challenge. However, their expression is very strong in primed plants after rechallenge. They are, therefore, specific readout genes for the primed state. Remarkably, mutation in solely priming-readout gene *AT5G36970* (encoding NDR1/HIN1-like protein 25) impaired both infection-induced defense priming and systemic acquired resistance suggesting a previously unknown critical role of this gene in the systemic plant immune response.

## Introduction

Upon localized leaf infection by necrogenic microbes or after chemical treatment, the entire plant foliage can become primed for enhanced defense (Conrath et al. 2002, 2006, 2015). Primed leaves respond more strongly to physical injury or pathogen attack and they frequently express full-body resistance to multiple diseases (Ross 1961; Ryals et al. 1996; Conrath et al. 2002, 2006, 2015). Different from the full activation of defense responses upon primary infection priming causes only low fitness costs (Van Hulten et al. 2006; Martinez-Medina et al. 2016). In addition, priming is hardly prone to pathogen adaptation (Conrath et al. 2015; Martinez-Medina et al. 2016). Therefore, triggering defense priming is promising for practical agronomic use and interesting as a paradigm for plant signal transduction as well (Beckers and Conrath 2007; Conrath et al. 2015).

Defense priming comprises enhanced levels in the plasma membrane of microbial pattern receptors and coreceptors (Tateda et al. 2014). They include protein kinase flagellin-sensing 2 (FLS2, recognizing the bacterial flagellin epitope flg22), brassinosteroid-insensitive 1-associated receptor kinase 1 (BAK1, a coreceptor of FLS2), and chitin elicitor receptor kinase 1 (CERK1, the chitin and peptidoglycan receptor or coreceptor, respectively) (Tateda et al. 2014). The enhanced presence of microbial pattern receptors and coreceptors in the plasma membrane of primed cells increases their responsiveness to microbes harboring flagellin, chitin, or peptidoglycan (Tateda et al. 2014). Consistent with the role of FLS2, BAK1, and CERK1 in activating mitogen-activated protein kinase (MPK) signaling relays (Asai et al. 2002; Bender and Zipfel 2023), priming likewise encompasses enhanced levels of dormant, but activable MPK3 and MPK6 molecules (Beckers et al. 2009). Because of the enhanced levels of MPK3 and MPK6 in primed cells, more of these enzymes are activated upon stimulation of the microbial-pattern receptors thus amplifying the transducing signal and, ultimately, leading to enhanced defense (Beckers et al. 2009).

In addition to the enhanced levels of microbial pattern receptors and activatable MPK3 and MPK6 molecules, priming for enhanced defense includes covalent modification of DNA and histones in the promoter of defense genes, such as those encoding WRKY transcription factors with a role in plant defense (Jaskiewicz et al. 2011; Luna et al. 2012). The modification of DNA and histones primes the affected gene for enhanced transcription after further stimulation (Conrath, 2011; Jaskiewicz et al., 2011; Luna et al., 2012). Together, the enhanced levels of microbial-pattern receptors and dormant MPKs as well as the mounting of gene-conditioning chromatin modifications provide a memory to the priming-inducing event in that they prime cells for the superinduction of defense responses by physical challenge or microbial attack, associated with development of stress tolerance and systemic acquired disease resistance (SAR) (Conrath et al. 2015).

Surprisingly, although defense priming received much attention both as a promising concept for plant protection and a paradigm for cellular signal transduction, the identity of genes that are specifically expressed during priming (referred to in this paper as priming-marker genes) or whose expression is stronger in primed than unprimed plants after challenge (referred to as priming-readout genes) remained largely unknown. This is particularly surprising for the intensively studied Arabidopsis (*Arabidopsis thaliana*) - Pca (*Pseudomonas cannabina* pv. alisalensis; formerly called *Pseudomonas syringae* pv. maculicola ES4326) interaction (Katagiri et al. 2002; Baltrus et al. 2011; Sarris et al. 2013). Knowing the identity of marker and readout genes for priming would not only equip the plant community with novel tools for the research into priming, but also help expanding the knowledge of the phenomenon and support its translation to agricultural practice, e.g., through identifying priming-inducing chemistry or by breeding for an enhanced sensitivity to be primed (Beckers and Conrath, 2007).

So far, we and others often used *AT1G62300* (*WRKY6*), *AT4G23550* (*WRKY29*), and *AT4G23810* (*WRKY53*) as readout genes to assess defense priming in Arabidopsis (Jaskiewicz et al. 2011; Luna et al. 2012; Baum et al. 2019). Monitoring their expression advanced the research into priming but the weight of these loci as priming-readout genes remained unclear. A genome-wide record of marker and readout genes for priming simply was missing. We recently used formaldehyde-assisted isolation of regulatory DNA elements (FAIRE) to provide a genome-wide map of regulatory DNA sites in the primed foliage of Arabidopsis plants with local Pca infection (Baum et al. 2019). Supplemental whole-transcriptome shotgun sequencing of mRNA transcripts from systemic leaves of primed and unprimed plants, both before and after physical challenge, disclosed all Arabidopsis genes with expression before (possible priming-marker genes) and enhanced expression after (possible priming-readout genes) rechallenge. So far, these datasets remained insufficiently explored and marker genes for individual immunological conditions (primed or unprimed both before and after challenge) unconfirmed. Here, we introduce genes that we validated as suitable marker or readout genes for defense priming in Arabidopsis. Based on *in-silico* analyses we also predict interaction networks and subcellular mapping of the proteins encoded by marker and readout genes for priming. We also demonstrate that mutation of solely priming-readout gene *AT5G36970* (encoding NDR1/HIN1-like protein 25; NHL25) attenuates Pca-induced priming for enhanced defense gene activation and impairs SAR.

## Results

### Spotting and validating marker genes for priming

To spot and confirm or refute marker and readout genes for defense priming in Arabidopsis, we thoroughly reevaluated Supplemental Dataset S1 of our previous publication by Baum et al. (2019). The dataset contains genes whose expression is activated or repressed in one or more of four immunological conditions, that is mock-inoculated on local leaves (control) or Pca-inoculated on local leaves (i.e., systemically primed) both before and after systemic challenge (Fig. 1). The dataset also comprises information about whether chromatin in the promoter of individual genes has been unchanged or more or less opened in any of the four immunological states, as determined by FAIRE (Baum et al. 2019, 2020).

**Figure 1.**
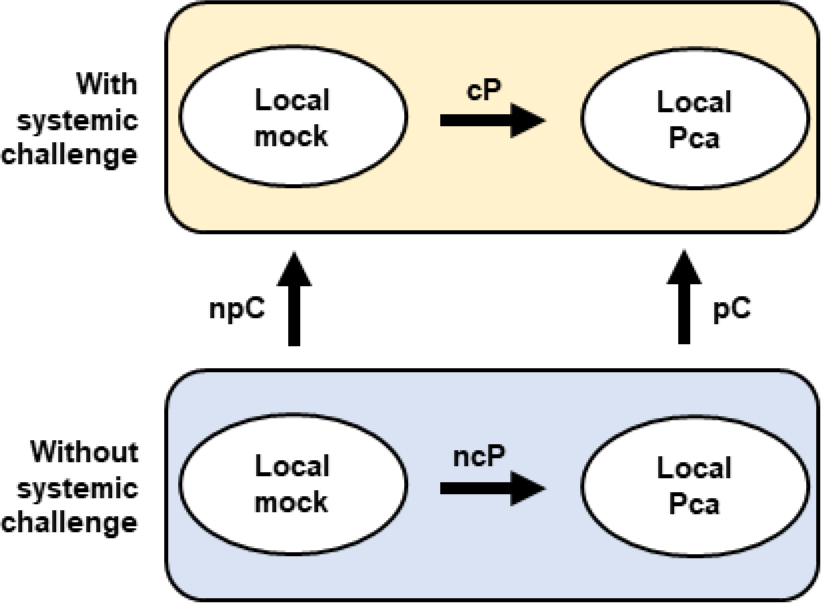
Immunological conditions used to identify marker and readout genes for defense priming in systemic leaves of Arabidopsis plants with local Pca infection. ncP, genes expressed during priming; npC, genes expressed after systemic challenge; pC, genes expressed after challenge in primed leaves; cP, genes expressed because of priming in systemic challenge condition.

We first used dataset ncP (Fig. 1) which should contain genes whose expression is activated during priming in infection-free systemic leaves of Arabidopsis plants with local Pca infection. We also considered dataset pC (Fig. 1) to spot those genes in the ncP dataset whose expression is reduced after systemic rechallenge in Pca-primed plants (as found in the pC dataset). Genes with high ncP and low pC values are most promising to be exclusive marker genes for defense priming. This is particularly true if they also have priming-associated open chromatin in the promoter, as inferred from the FAIRE dataset of Baum et al. (2019).

We *in silico* determined the top ten ncP genes with high FAIRE and low pC value (Tab. 1). We *in silico* also identified the top cP genes with high FAIRE value (Tab. 1). Then, we in the lab reassessed the expression of these genes in any of the four immunological conditions (control, primed, challenged, and primed followed by challenge). Of the genes analyzed, *AT1G76960* (encoding a protein with unknown function) (Fig. 2A), *AT3G51860* (vacuolar Ca^2+^/H^+^ antiporter CAX3) (Fig. 2B), *AT3G45860* (cysteine-rich receptor-like protein kinase CRK4) (Fig. 2C), and *AT5G64190* (neuronal PAS-domain protein with unknown function) (Fig. 2D), the latter with low overall expression, are particularly expressed during Pca-induced priming in infection-free systemic leaves but not, or to a lesser extent, in control or challenged leaves, or in primed and subsequently challenged leaves (Fig. 2). Expression of these genes thus solely marks the primed state of enhanced defense readiness.

**Figure 2.**
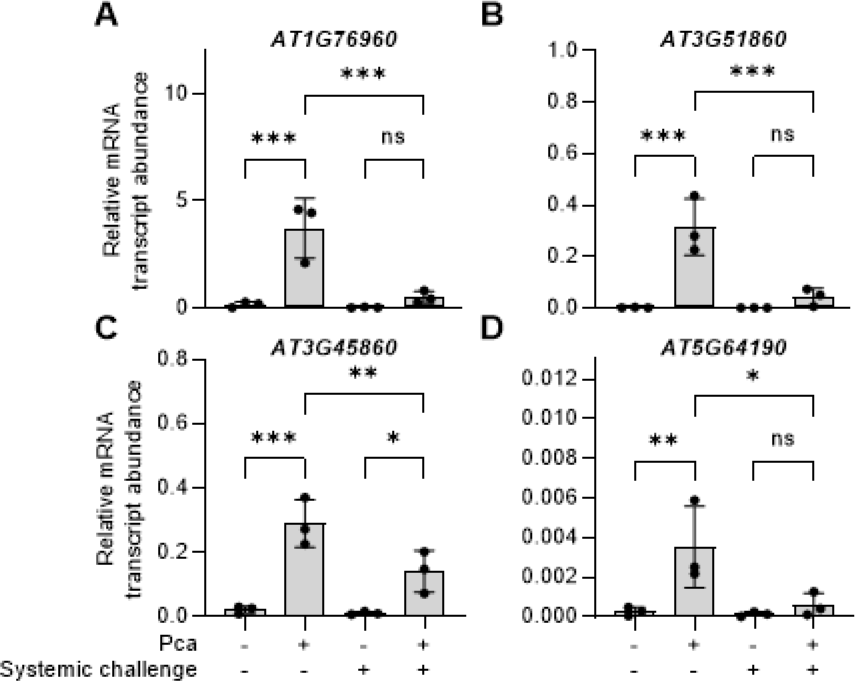
Expression of genes *AT1G76960* (**A**), *AT3G51860* (*CAX3*; **B**), *AT3G45860* (*CRK4*; **C**) and *AT5G64190* (**D**) is particularly activated during priming. Five-week-old Arabidopsis plants were infiltrated on three leaves with MgCl_2_ (mock inoculation; - Pca) or a Pca suspension in MgCl_2_ (+ Pca). Three days later, untreated leaves of both sets of plant were left untreated (-systemic challenge) or challenged by the infiltration of water (+ systemic challenge). Three h later, the systemic leaves were harvested and analyzed for the expression of specified genes. Relative mRNA transcript abundance was determined by RT-qPCR and normalized to the expression of *ACTIN2*. Shown are the values and SD of three independent experiments, each with two plants. Statistical significance was determined using Ordinary one-way ANOVA. ***P < 0.001; **P < 0.01; *P < 0.05; ns, not significant.

**Table 1.**
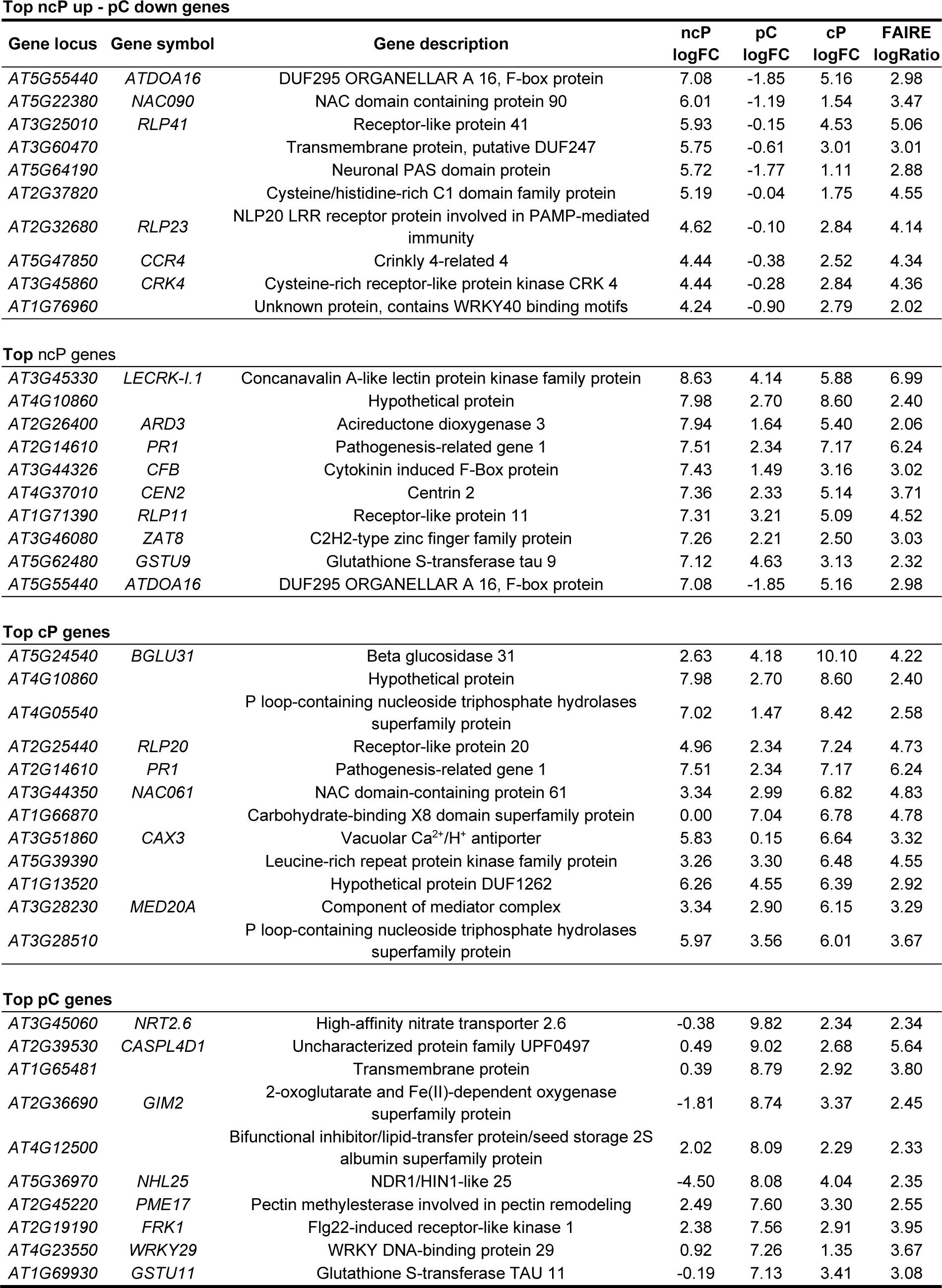
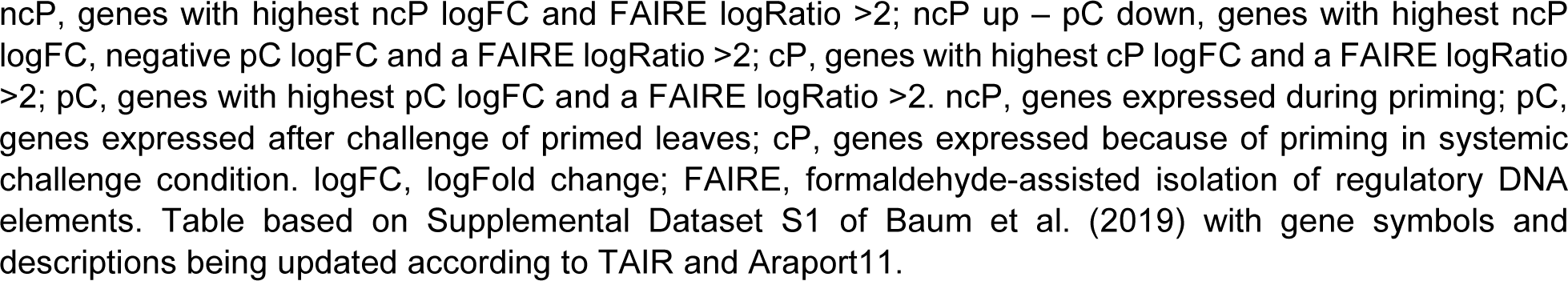
Top candidates for marker and readout genes for defense priming.

Different from the above-mentioned genes, *AT2G14610* (encoding pathogenesis-related protein PR1) (Fig. 3A), *AT2G32680* (receptor-like protein RLP23) (Fig. 3B), *AT3G25010* (RLP41) (Fig. 3C), *AT5G47850* (crinkly 4-related 4, CCR4) (Fig. 3D), *AT4G05540* (for a P loop-containing nucleoside triphosphate hydrolases superfamily protein) (Fig. 3E), and *AT1G66870* (carbohydrate-binding X8-domain superfamily protein) (Fig. 3F) are activated, to about equal extent, in primed leaves before and after rechallenge (Fig. 3). They can also be used as marker genes for priming as their expression is activated in primed condition and not much changed upon rechallenge (Fig. 3).

**Figure 3.**
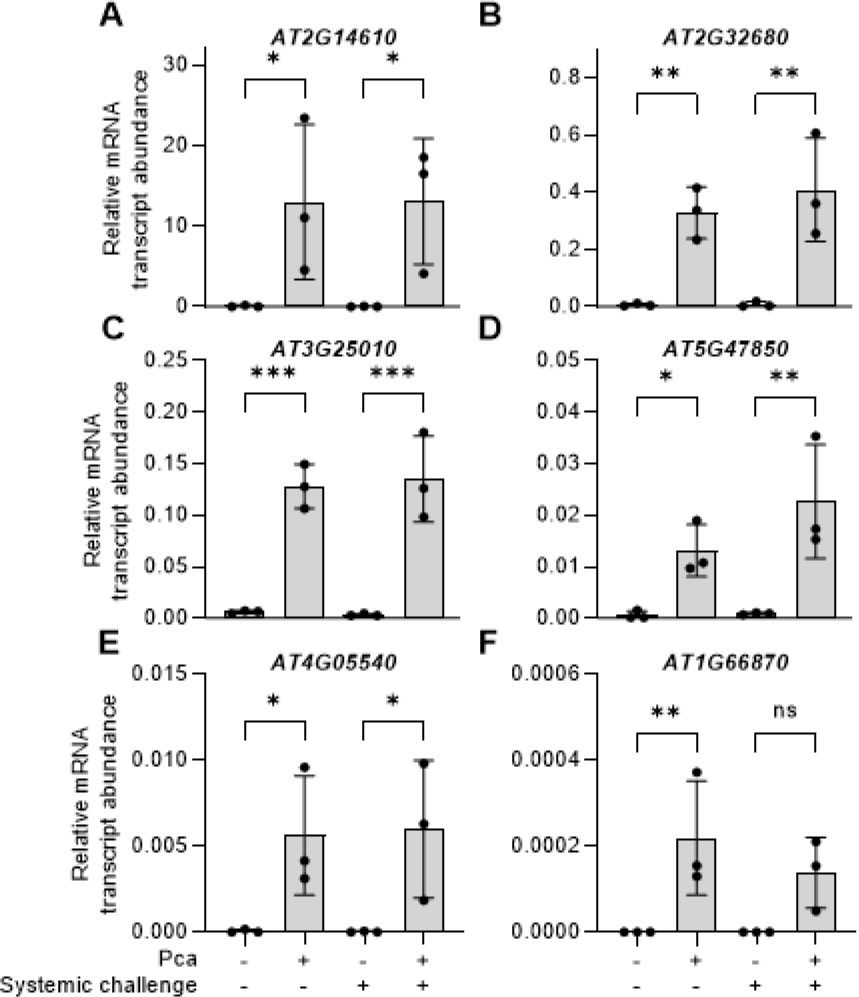
Expression of genes *AT2G14610* (*PR1*; **A**), *AT2G32680* (*RLP23*; **B**), *AT3G25010* (*RLP41*; **C**), *AT5G47850* (*CCR4*; **D**), *AT4G05540* (**E**), and *AT1G66870* (**F**) is associated with defense priming. Experimental setup, data and statistical analyses were performed as in Figure 2. ***P < 0.001; **P < 0.01; *P < 0.05; ns, not significant.

### Spotting and validating readout genes for priming

We next aimed at identifying and possibly verifying genes one can use as specific readout genes for defense priming in Arabidopsis. To do so, we *in silico* reassessed our pC, cP, ncP, and FAIRE datasets (Fig. 1; Tab. 1; Baum et al. 2019). We picked eight genes with high expression after systemic rechallenge of primed leaves (mainly in the pC dataset) and a priming-associated open chromatin (FAIRE) value of >2 (Baum et al. 2019). We in the analysis also included previously used priming-readout genes *AT1G62300* (transcription factor gene *WRKY6*; #84 in the pC gene list of Baum et al. (2019) and *AT4G23810* (*WRKY53*; #232 in their pC list of genes (Supplemental Table S1, Baum et al. 2019) to assess their weight as readout genes for the primed state (Jaskiewicz et al. 2011; Luna et al. 2012).

Genes *AT2G39530* (encoding Casparian strip domain-like protein [CASPL]4D1) (Fig. 4A), *AT2G19190* (flg22-induced receptor-like kinase FRK1) (Fig. 4B), *AT3G28510* (P loop-containing nucleoside triphosphate hydrolases superfamily protein) (Fig. 4C), *AT4G12500* (bifunctional inhibitor/lipid-transfer protein/seed storage 2S albumin superfamily protein) (Fig. 4D), *AT1G62300* (WRKY6) (Fig. 4E), *AT4G23810* (WRKY53) (Fig. 4F), *AT2G45220* (pectin methyl esterase PME17) (Fig. 4G), *AT4G23550* (WRKY29) (Fig. 4H), *AT3G45060* (encoding high-affinity nitrate transporter [NRT]2.6) (Fig. 4I), and *AT1G71390* (for receptor-like protein 11) (Fig. 4J) are not, or hardly expressed during priming or after challenge. But they are strongly expressed in primed leaves following challenge (Fig. 4). Those genes thus classify as specific readout genes of priming.

**Figure 4.**
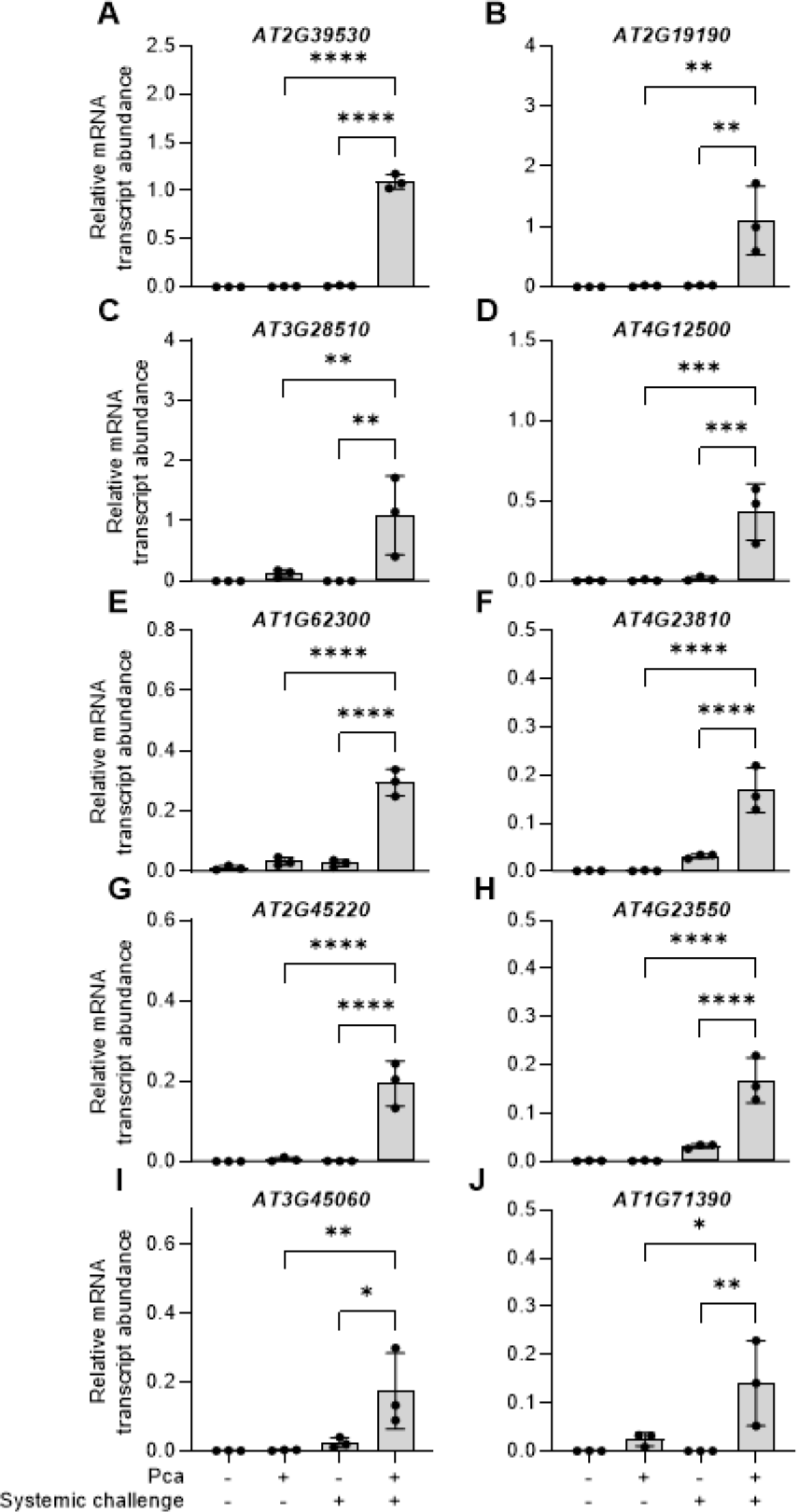
Readout genes of defense priming in Arabidopsis. Genes AT2G39530 (CASPL4D1, **A**), AT2G19190 (FRK1, **B**), AT3G28510 (**C**), AT4G12500 (**D**), AT1G62300 (WRKY6, **E**), AT4G23810 (WRKY53, **F**), AT2G45220 (PME17, **G**), AT4G23550 (WRKY29, **H**), AT3G45060 (NRT2.6, **I**), and AT1G71390 (RLP11, **J**) are particularly expressed in primed leaves when these have been challenged. Experimental setup, data and statistical analyses were done as in Figure 2. ****P < 0.0001; ***P < 0.001; **P < 0.01; *P < 0.05.

### Analyzing the interaction of proteins encoded by marker and readout genes for priming

Disclosing interaction networks of genes and proteins can help understand biological processes at the systems level. To provide insight into defense priming in Arabidopsis at this level we used the Search Tool for the Retrieval of Interacting Genes/Proteins (STRING) (Szklarczyk et al. 2021) (https://string-db.org) and disclosed possible interaction networks of genes and proteins during defense priming in this plant. The STRING analysis for validated specific marker (Fig. 2) and readout genes (Fig. 4) for priming predicted, with high confidence (STRING value ≥ 0.7), an equally-intense interaction network of priming-readout proteins AT2G19190 (FRK1), AT1G62300 (WRKY6), and AT4G23810 (WRKY53) (Fig. 5A). Disclosure of the protein interaction network was based on co-expression of the encoding genes (Supplemental Dataset S1). With medium confidence (STRING value ≥ 0.4) AT2G19190 (FRK1) also seems to interact with AT4G23550 (WRKY29) (Fig. 5A). Intriguingly, we found clues to an intense interaction of priming-marker protein AT3G45860 (CRK4) with the P loop-containing nucleoside triphosphate hydrolases superfamily protein encoded by priming-readout gene *AT3G28510* (Fig. 5A). Not only were the genes encoding the two proteins found to be co-expressed but also seem the proteins encoded by orthologous genes in man, mouse and the eelworm *Caenorhabditis elegans* to interact with each other. These findings strongly suggest that CRK4 and the *AT3G28510*-encoded P loop-containing nucleoside triphosphate hydrolases superfamily protein interact in Arabidopsis too.

**Figure 5.**
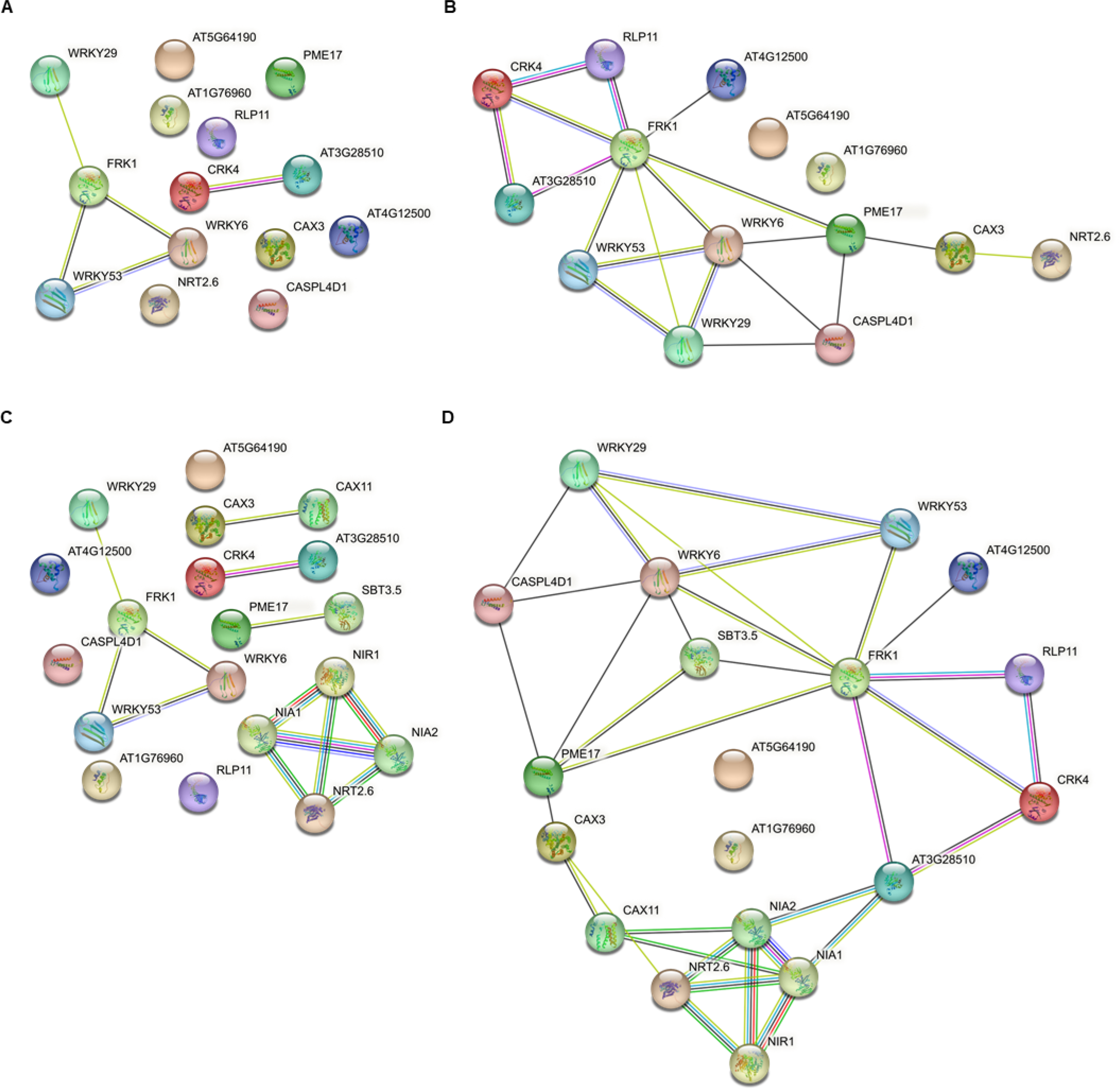
Associations of proteins encoded by marker or readout genes of defense priming. **A), B),** Direct interactions. **A),** Medium confidence (STRING value ≥ 0.4) or **B),** low confidence (STRING value ≥ 0.15). **C)**, **D),** Interactions after allowing additional nodes. **C),** Medium confidence and **D),** low confidence. Nodes represent proteins. Unfilled nodes represent proteins with unknown 3D structure. Filled nodes represent proteins with known or predicted 3D structure. Edges represent protein-protein associations. Known interactions (in cyan blue from curated databases, in purple experimentally determined), predicted interactions (green, gene neighborhood; red, gene fusions; blue, protein co-occurrence); Others (yellow, text mining; black, co-expression; light blue, protein homology). Figures were drawn using STRING database.

In addition to AT1G62300 (WRKY6), AT4G23810 (WRKY53), and AT4G23550 (WRKY29), AT2G19190 (FRK1), with low-to-medium confidence, also seems to interact with priming-readout protein AT1G71390 (RLP11), the P loop-containing nucleoside triphosphate hydrolases superfamily protein encoded by priming-readout gene *AT3G28510*, and priming-marker protein AT3G45860 (CRK4) (Fig. 5B). AT2G19190 (FRK1) and AT1G62300 (WRKY6), with medium confidence, seem to be central nodes in an interaction network that contains all the in this paper validated priming-marker and readout proteins, except the neuronal PAS-domain protein encoded by *AT5G64190* and the protein with unknown function encoded by *AT1G76960* (Fig. 5B).

When we lowered the stringency during node identification (Fig. 5C and D) our analysis with medium confidence disclosed an interaction of priming-marker protein AT3G51860 (CAX3) with AT1G08960 (CAX11), of AT2G45220 (PME17) with AT1G32940 (subtilase-family protein SBT3.5) (Fig. 5C), and of high-affinity nitrate transporter NRT2.6 (AT3G45060) with nitrate reductases NIA1 (AT1G77760) and NIA2 (AT1G37130), and nitrite reductase NIR1 (AT2G15620) (Fig. 5C). The latter four proteins, which have all been associated with nitrogen metabolism (Dechorgnat et al. 2012), are highly likely to interact. Based on gene neighborhood and co-expression, CAX11 (AT1G08960) with low confidence seems to interact with NIA1(AT1G77760) and NIA2 (AT1G37130) (Fig. 5D).

In sum, our STRING interaction network analysis disclosed that the genes *AT4G23810* (encoding WRKY53) and *AT2G19190* (encoding FRK1) often seemed to be co-expressed (Fig. 5A-D). Especially AT2G19190 (FRK1) seems to be a central player in the priming network of proteins. In fact, many interactions that we predicted for the FRK1 protein have been experimentally demonstrated in pull-down assays or by solid phase array analysis (Tab. 2; Smakowska-Luzan et al. 2018; Mott et al. 2019). Interestingly, for the neuronal PAS-domain protein encoded by *AT5G64190* putative interaction partners were previously predicted only with low confidence (0.15), based on *AT5G64190*’s co-expression with *AT4G19420* (for a pectin acetylesterase-family protein) and *AT3G03870* (encoding a protein with unknown function). Remarkably, until now co-expression and interaction has barely been described for most of the here spotted marker and readout genes or proteins, respectively.

**Table 2.**
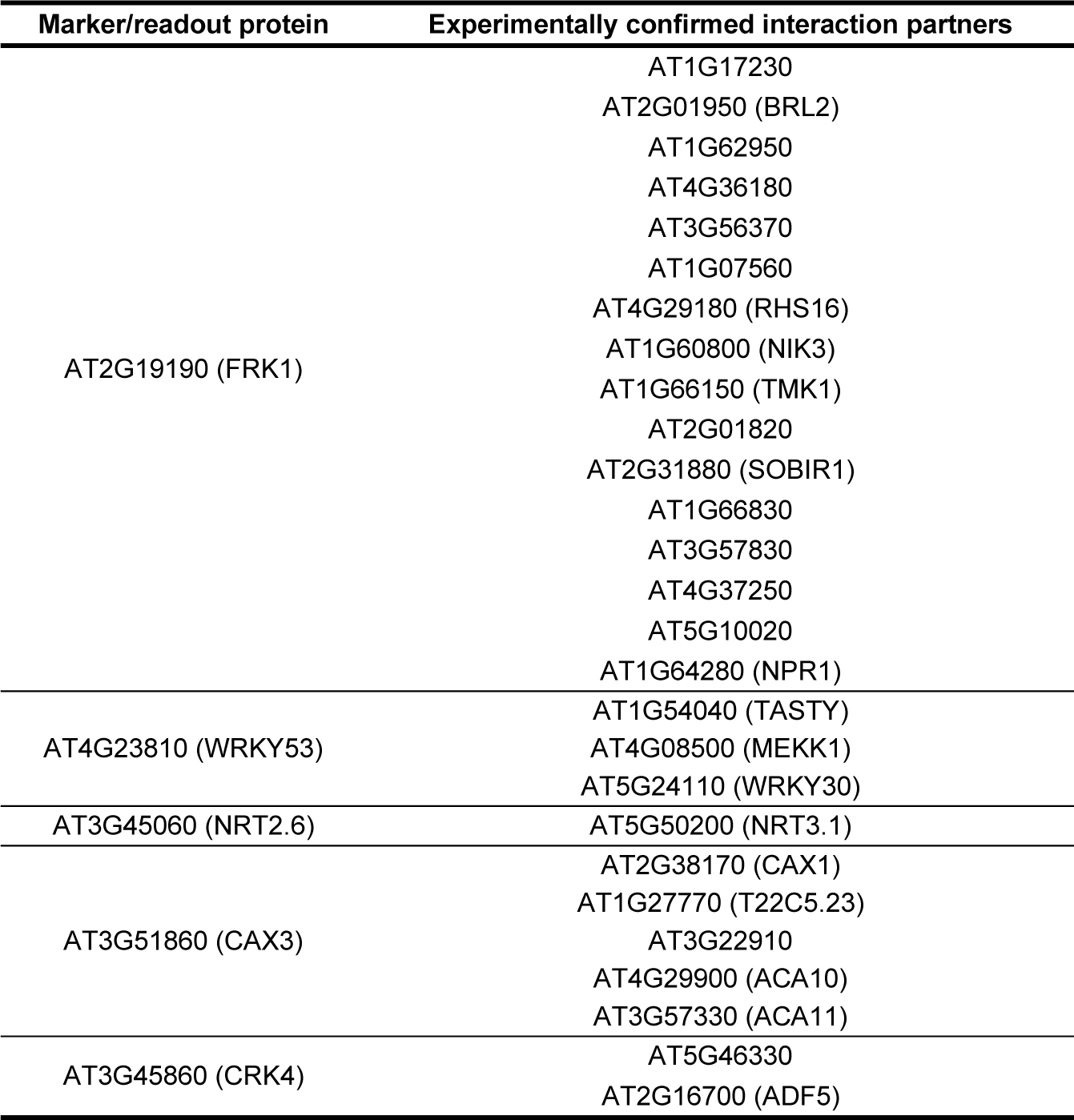
Experimentally demonstrated interactions of proteins encoded by marker and readout genes of priming (high-confidence; STRING value ≥0.7).

### Subcellular localization of marker and readout proteins for priming and their co-accumulating partners

To get even more insight into defense priming, we further analyzed the results of our STRING analysis based on the co-expression of genes (threshold STRING co-expression score 0.4). We used the Subcellular Localization database for Arabidopsis proteins (SUBA, version 5) (Hooper et al. 2022) (http://suba.live/) to determine the localization of interesting proteins within the cell. We did so by considering only candidates with a connection to the 14 genes in Figures 2 and 4 and a high cut-off (≥ 0.5 stringency). We drew a subcellular priming map of proteins (Fig. 6). As shown in Figure 6, most of the proteins that we assume to interact with the in this work identified marker and readout proteins for priming are localized to the plasma membrane (15 in total), nucleus (13), or extracellular space (6). Three of them are localized to the mitochondria, and one to each, the cytosol and peroxisome (Fig. 6). The distribution of marker and readout proteins for priming and their interacting proteins in at least five cellular compartments indicates physiological complexity for defense priming. In addition, the predominance in the plasma membrane and extracellular space of these proteins substantiates their important defensive role in the apoplast/extracellular space.

**Figure 6.**
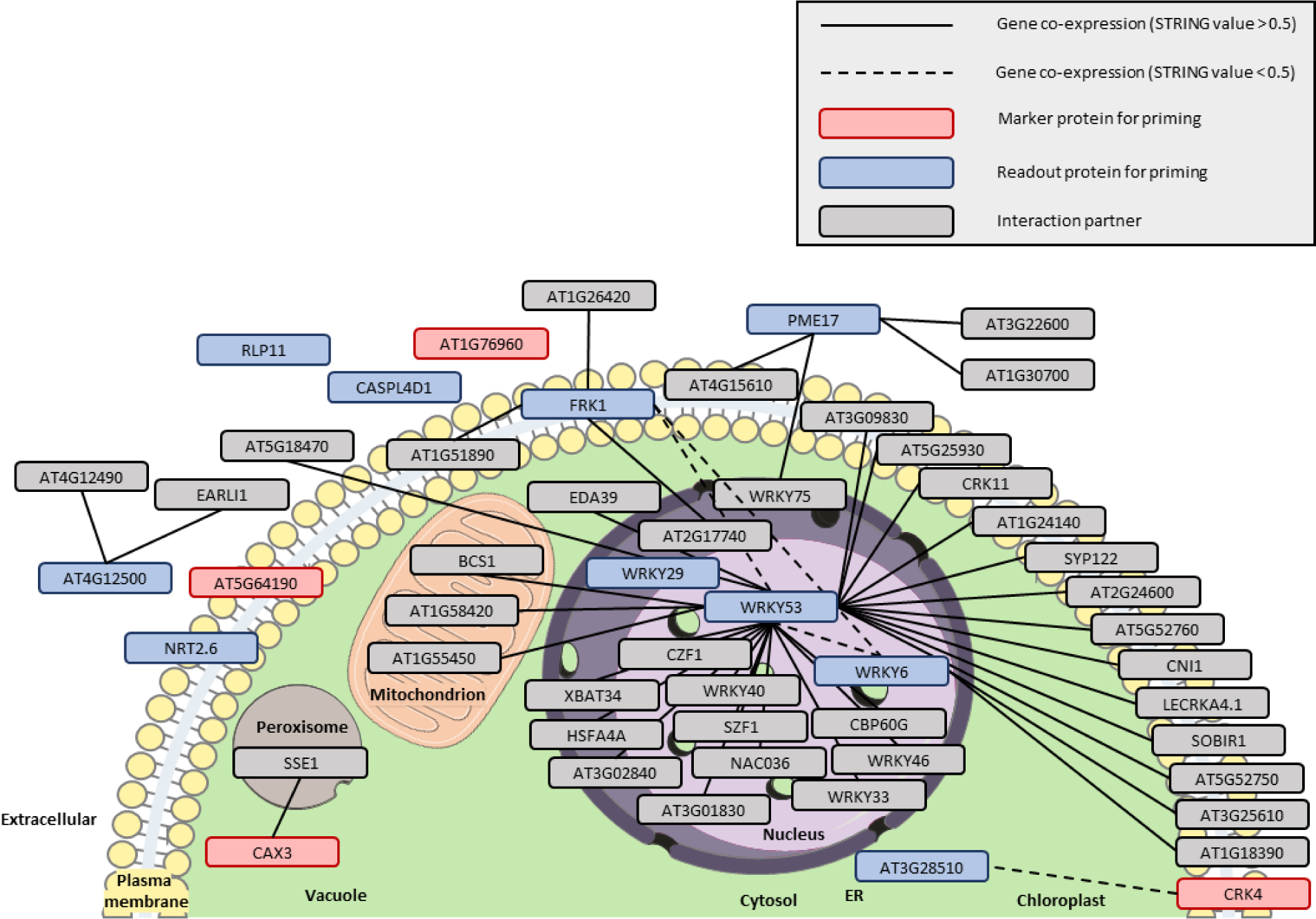
Subcellular localization of marker and readout proteins for priming and their supposed interaction partners. We used SUBA to determine the subcellular localization of marker and readout proteins for priming and their presumed-interacting protein partners. To keep clarity, we raised the STRING value to ≥ 0.5 when analyzing the co-expression of genes other than the identified marker or readout genes for priming (Figs. 2 and 4). ER, endoplasmic reticulum. Parts of the figure were drawn by using pictures from Servier Medical Art. Servier Medical Art by Servier is licensed under a Creative Commons Attribution 3.0 Unported License (https://creativecommons.org/licenses/by/3.0/).

### Mutation of priming-readout gene *AT5G36970* (*NHL25*) attenuates priming and impedes SAR

In Arabidopsis expression of *AT5G36970* (encoding NDR1/HIN1-like protein NHL25) is induced during incompatible interactions with pathogens (Varet et al. 2002). In addition, *AT5G36970* (*NHL25*) expression, at least in part, depends on salicylic acid (Varet et al. 2002) which primes plants for enhanced defense (Thulke and Conrath 1998; Katz et al. 2002). Thus, *NHL25*, which is #6 in the pC dataset (Table 1) could be important for defense priming and SAR. We have taken this into account, and we found that the *NHL25* gene is hardly expressed in untreated leaves, during priming or after challenge (Fig. 7A). However, *NHL25* is expressed in primed leaves after they have been challenged (Fig. 7A). Thus, *NHL25* qualifies as a bona-fide readout gene for defense priming in Arabidopsis. Surprisingly, although defense priming seems to be a complex physiological phenomenon (Fig. 6) mutation of merely *NHL25* impaired the Pca-induced systemic expression of priming-marker genes *AT1G76960* (for a protein with unknown function), *AT3G51860* (*CAX3*), and *AT3G45860* (*CRK4*) (Fig. 7B-D) as well as of priming-readout genes *AT2G39530* (*CASPL4D1*), *AT2G19190* (*FRK1*), and *AT3G28510* (for a P loop-containing nucleoside triphosphate hydrolases superfamily protein) (Fig. 7E-G). Moreover, mutation of *NHL25* impedes the development of SAR to Pca in two independent *nhl25* and the priming/SAR-negative *npr1* mutant (Kohler et al. 2002) (Fig. 7H; Supplemental Fig. S1). Remarkably, the two independent *nhl25* mutants, different than *npr1*, do not express enhanced basal susceptibility to Pca (Fig. 7H; Supplemental Fig. S1). Together the findings in Figure 7B-H disclose *NHL25* as a previously unknown key gene in the primed SAR response of Arabidopsis.

**Figure 7.**
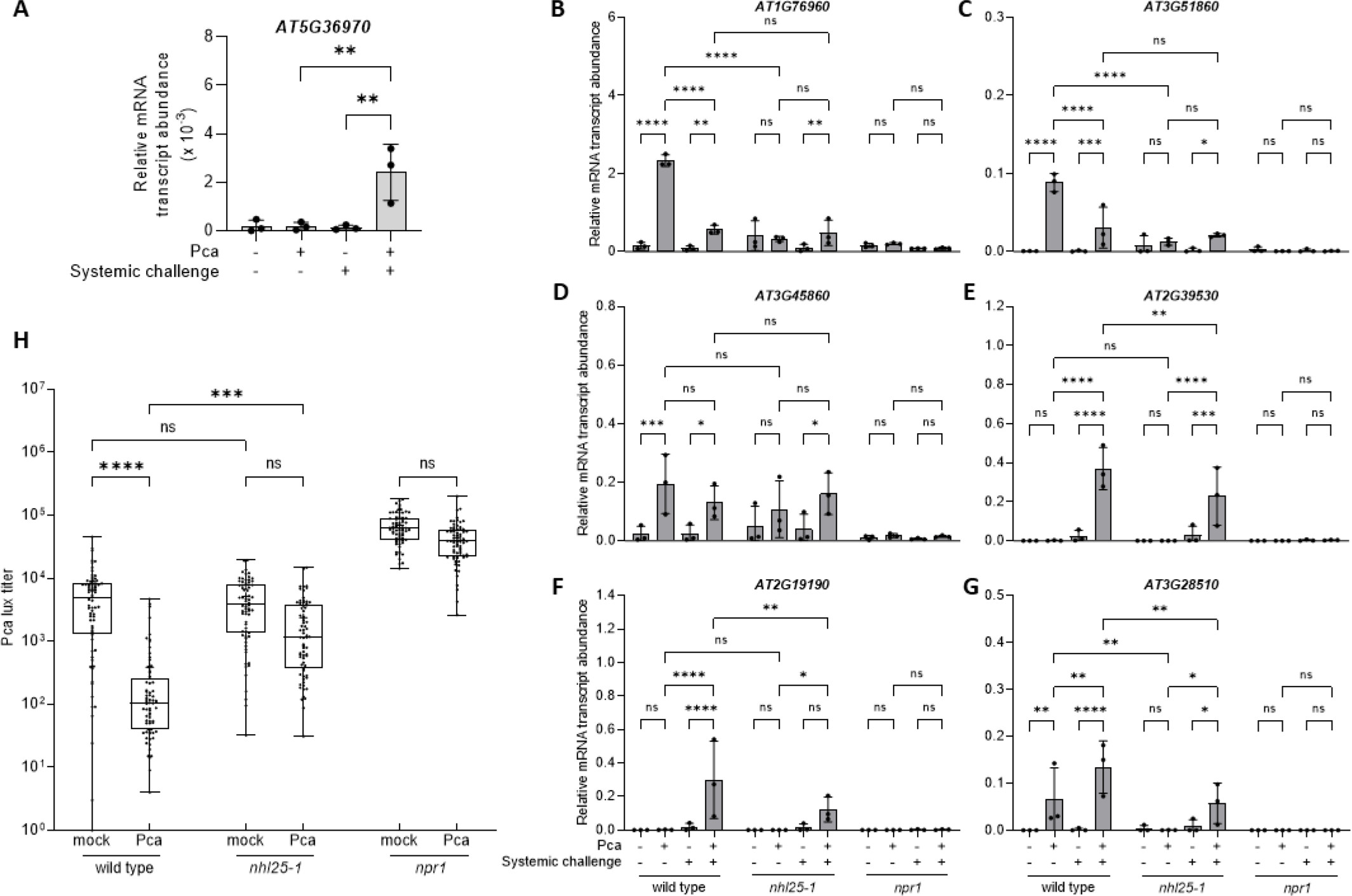
*NHL25* is a priming-readout gene whose mutation impairs defense priming and SAR. **A).** *NHL25* is a readout gene for priming. Five-week-old wild-type plants were mock-inoculated (-Pca) or infected with Pca (+) on three leaves. Three d later, distal leaves were left untreated (-systemic challenge) or infiltrated with water (+ systemic challenge). Three h later, the systemic leaves were harvested and analyzed for the expression of the *NHL25* gene (normalized to *ACTIN2*). **B) - G).** Priming is attenuated or absent in the *nhl25-1* (SALK_113216) and *npr1* mutant. Five-week-old Arabidopsis plants were mock-inoculated (-Pca) or infected with Pca (+) on three leaves. Three days later, systemic leaves were left untreated (-systemic challenge) or challenged by the infiltration of water (+ systemic challenge). Three h later, untreated or infiltrated systemic leaves were harvested and analyzed for expression of the indicated marker **B) - D)** and readout **E) - G)** genes of priming. Relative mRNA transcript abundance was determined by RT-qPCR and normalized to *ACTIN2*. **H).** SAR is absent in the *nhl25-1* and *npr1* mutant. Five-week-old plants were mock-inoculated or infected with Pca on three leaves. Three days later, uninoculated systemic leaves were inoculated with Pca lux. After another 3 d, the titer of Pca lux was determined by measuring the luminescence in discs taken from systemic Pca lux-inoculated leaves. For **A) - C)**, data and SD of three independent experiments each with two plants are shown. For **D)**, data derived from three independent experiments each with eight biological replicates consisting of three leaves from an appropriately treated plant. Statistical significance was tested with Ordinary one-way ANOVA **A)** - **C**) or with Kruskal-Wallis test (**D**). ****P < 0.0001; ***P < 0.001; **P < 0.01; *P < 0.05; ns, not significant.

## Discussion

Marker genes are useful to explore the physiological condition of a cell, tissue, or organism in a given condition, e.g., in response to biotic or abiotic stress. Defense priming is a state of the enhanced defense readiness in immunostimulated cells (Conrath et al. 2002, 2006, 2015). The identity of marker and readout genes for defense priming remained largely unknown. Knowing them will advance the research into priming and facilitate its application in agriculture (Conrath et al. 2015; Beckers and Conrath, 2007). Based on the in this paper disclosed pattern and level of expression, we recommend using *AT1G76960* (for a protein with unknown function), *AT3G51860* (CAX3), and *AT3G45860* (CRK4) as exclusive marker genes for defense priming in the Arabidopsis-Pca interaction (Fig. 2). *AT2G14610* (PR1) because of its very strong expression (Fig. 3A) is also recommended for determining the presence of priming in this plant. However, the expression of *PR1* is not exclusive to the ‘only primed’ state. *AT2G39530* (*CASPL4D1*), *AT2G19190* (*FRK1*), and *AT3G28510* (for the P loop-containing nucleoside triphosphate hydrolases superfamily protein) should be used as priming-readout genes (Fig. 4). In principle, the other in this work identified marker (Figs. 2D, 3) and readout (Fig. 4D-J) genes for priming can also be used for the research into defense priming in Arabidopsis. However, their overall expression level is lower than of the genes recommended above.

The specific priming-marker gene *AT3G51860* encodes vacuolar Ca^2+^/H^+^ antiporter CAX3 that forms a complex with AT1G16380 (CAX1) (Fig. 5C), another Ca^2+^/H^+^ antiporter in the tonoplast (Berardini et al. 2015). Both proteins contribute to Ca^2+^ homeostasis by catalyzing Ca^2+^ transport from the cytosol into the vacuole by exploiting the proton gradient across the tonoplast (Cho et al. 2012). Ca^2+^ serves as second messenger (White and Broadley 2003) and activates defense responses in plants (Zhang et al. 2014). In unstimulated cells, its concentration is only low (∼100nM) (White and Broadley 2003; Zhang et al. 2014). The Ca^2+^ concentration rapidly rises upon microbial pattern recognition, mainly by influx from the apoplast through specific Ca^2+^ channels (Zhang et al. 2014). The resulting rise in cytosolic Ca^2+^ is sensed by Ca^2+^-binding proteins (e.g., calmodulin, Ca^2+^-dependent protein kinases, calcineurin B-like proteins) that translate the Ca^2+^ signal to cellular responses that, e.g., help fight off infection (Zhang et al. 2014). Because high cytosolic Ca^2+^ concentrations are toxic, excessive Ca^2+^ ions need to be removed from the cytosol (White and Broadley 2003). This is catalyzed by Ca^2+^-ATPases and Ca^2+^/H^+^ antiporters, such as CAX1 and CAX3 (Hirschi 2001). Thus, the priming-linked expression of *AT3G51860* (*CAX3*) might be in anticipation of challenge-induced threatening rises in cytosolic Ca^2+^ concentration thus avoiding exaggerating intracellular Ca^2+^ increase as part of an anti-inflammatory response.

Priming-marker gene *AT3G45860* encodes CRK4 that in the plasma membrane associates with flg22 receptor FLS2 and, in a possible interaction with CRK6 and CRK36 contributes to the priming for an enhanced flg22-induced oxidative burst and the defense of pathogenic Pseudomonads (Yeh et al. 2015). Just like MPK3, MPK6, and FRK1, CRK4 during priming could accumulate in its inactive, yet activable, form (Beckers et al. 2009). Upon perception of microbial patterns, such as flg22, more CRK4 molecules could be activated in primed compared to unprimed cells, thereby amplifying the transducing signal and, eventually, cause more robust defense (Beckers et al. 2009). However, whether CRK4 has this function remains to be seen.

*AT2G14610* encodes PR1 that strongly accumulates after infection by many pathogens and upon treatment with certain chemicals, including salicylic acid and its biologically active derivatives (Ward et al. 1991; Conrath et al. 1995). PR1 has been used as a molecular marker for SAR in different plant species including Arabidopsis for a long time (Van Loon et al. 2006). The PR1 protein binds sterols and causes cellular leakage (Gamir et al. 2017). By doing so it exerts antimicrobial activity that has been demonstrated both *in vitro* (Niderman et al. 1995) and in transgenic plants overexpressing *PR1* (Alexander et al. 1993). Arabidopsis PR1 proteins are secreted to the extracellular space (Van Loon et al. 2006) where they can directly fight pathogens.

Priming-readout gene *AT2G39530* encodes CASPL4D1 (Berardini et al. 2015). The membrane protein is localized to the chloroplast and plasma membrane and has been associated with the defense response of Arabidopsis to Pseudomonas (Mohr and Cahill 2007). However, CASPL4D1’s mode of action and role in plant immunity remain unclear. *AT2G19190*, in contrast, encodes FRK1 (Po-Wen et al. 2013; Berardini et al. 2015) that seems to be a central player in the priming network of proteins (Fig. 5; Table 2). Based on the expression of *FRK1* in the immunological conditions analyzed, the FRK1 protein seems not to accumulate during priming (Fig. 4B). Rather, *FRK1* expression is strongly activated when primed plants are being challenged (Fig. 4B) that is in a state of greatest distress.

Interestingly, *FRK1* is a reported target of transcription factor WRKY6 (Robatzek and Somssich 2002). Our STRING analysis disclosed that FRK1 most likely also interacts with WRKY53 (AT4G23810) and WRKY29 (AT4G23550) (Fig. 5A). We previously used *WRKY6* (*AT1G62300*), *WRKY29* (*AT4G23550*), and *WRKY53* (*AT4G23810*) as priming-readout genes (Jaskiewicz et al. 2011) but their weight to serve as such remained elusive. *WRKY29* is #9 in our top pC list (Tab. 1) and *WRKY6* and *WRKY53* do also have reasonably high pC and FAIRE, and rather low cP values (Supplemental Table S1, Baum et al. 2019) assigning them to the top ten priming-readout genes in their investigation (Fig. 4). *WRKY6*, *WRKY29*, and *WRKY53* belong to a family of >70 loci encoding transcription factors with a regulatory role in plant immunity (Wang et al. 2006; Jaskiewicz et al. 2011; Tsuda and Somssich, 2015). Priming of the three genes for enhanced transcription involves modification to histones in the promoter of the genes, notably trimethylation of histone H3K4 and acetylation of histone H3K9, as well as of histone H4K5, H4K8, and H4K12 (Jaskiewicz et al. 2011). These histone modifications provide docking sites for chromatin-regulatory proteins whose binding leads to local nucleosome eviction associated with the formation of nucleosome-free DNA (open chromatin) which is a hallmark of primed gene promoters (Baum et al. 2019; 2020; Giresi and Lieb 2009).

The *AT3G28510*-encoded P loop-containing nucleoside triphosphate hydrolases superfamily priming-readout protein (Berardini et al. 2015) most likely interacts with priming-marker protein AT3G45860 (CRK4) (Fig. 5A). This *in-silico* finding discloses a direct interaction of a priming-marker protein with a priming-readout polypeptide. In the incompatible interaction of Arabidopsis with *P. syringae* expressing the bacterial effector protein *avrRpt2*, the expression of *AT3G28510* (encoding the P loop-containing nucleoside triphosphate hydrolases superfamily protein) requires functional NDR1 (AT3G20600) (Sato et al. 2007) which is an important protein in the disease resistance response of Arabidopsis to bacterial and fungal pathogens (Century et al. 1995). The co-expression of NDR1-specific priming-readout genes *AT3G28510* and *AT2G19190 (FRK1)* (Fig. 5B) suggests involvement of the P loop-containing nucleoside triphosphate hydrolases superfamily protein AT3G28510 in the Arabidopsis MAMP-response pathway. This further qualifies *AT3G28510* as a robust priming-readout gene. The role in priming of the newly disclosed, likely interaction of AT3G28510 with priming-marker polypeptide CRK4 (Fig. 5A) remains to be experimentally confirmed. The same is true for the interaction of AT3G51860 (CAX3) with AT1G08960 (CAX11) (Fig. 5C and D) and the interplay of AT2G19190 (FRK1) with AT1G71390 (RLP11) (Fig. 5B) and the P loop-containing nucleoside triphosphate hydrolases superfamily protein AT3G28510 with AT3G45860 (CRK4) (Fig. 5C). Together, it seems that priming involves enhanced levels of multiple receptors with a role in plant defense that directly or indirectly, for instance via MPK cascades and other signal amplifying routes, target priming-readout genes to enable the primed immune response. The subcellular localization of in this work validated marker proteins together with the strong co-expression of genes encoding their interacting partner proteins (Fig. 5; Table 2) provides insight into the intrinsic interaction network between different cellular compartments.

We find the suggested involvement of nitrate metabolism proteins AT1G08090 (NRT2.6), AT1G77760 (NIA1), AT1G37130 (NIA2), and AT2G15620 (NIR1) (Fig. 5C) in defense priming intriguing. While AT1G08090 (NRT2.1), AT1G08100 (NRT2.2), AT5G60770 (NRT2.4), and AT5G14570 (NRT2.7) are proven nitrate transporters whose genes are being induced at low nitrogen levels, *AT1G08090* (*NRT2.6*) expression is rather activated at high nitrogen levels (Dechorgnat et al. 2012). In addition, a *nrt2.6* mutant did not show a nitrate-related phenotype (Dechorgnat et al. 2012). Consistently, even strong *NRT2.6* overexpression did not rescue the nitrate-uptake defect of a *nrt2.1-nrt2.2* double mutant (Dechorgnat et al. 2012). These findings suggested that Arabidopsis NRT2.6 may have other, or additional roles than transporting nitrate. Interestingly, *NRT2.6* expression was activated in Arabidopsis upon *Erwinia carotovora* infection and, consistently, plants with reduced *NRT2.6* expression were more susceptible to this pathogen (Dechorgnat et al. 2012). The authors suggested a link between *NRT2.6* expression and the fight of Arabidopsis against *E. carotovora* via the accumulation of reactive oxygen species (Dechorgnat et al. 2012). Our here reported findings support such a role of *NRT2.6* in pathogen defense (Fig. 4I). The obvious link between nitrogen metabolism and defense priming could be due to nitrate reductase-mediated release of nitric oxide which occurs at exaggerate nitrate reductase activity (Desikan et al. 2002; Yamamoto-Katou et al. 2006). During optimal nitrogen assimilation, cytoplasmic nitrate reductase catalyzes the reduction of nitrate to nitrite. The latter compound is transported to the chloroplast where it becomes reduced to NH_4_^+^ by nitrite reductase. Thus, AT1G77760 (NIA1) and AT1G37130 (NIA2) usually do not come together with AT2G15620 (NIR1). However, the cytoplasmic nitrite concentration can increase to high levels in the absence of an electrochemical gradient across the chloroplast envelope. This occurs, for example, when photosynthetic electron transport is impaired because of an attack by necrogenic pathogens.

We were surprised to find that mutation in merely *AT5G36970* (*NHL25*) impaired Pca-induced defense priming and SAR in Arabidopsis (Fig. 7B-H). The finding points to an important role of NHL25 in the two defense responses. Consistently *NHL25* is hardly induced in interactions of Arabidopsis with compatible Pseudomonads (Fig. 7A) (Varet et al. 2002). However, *NHL25* expression is strong during interactions of Arabidopsis with *P*. *syringae* pv. *tomato* DC3000 carrying avirulence gene *avrRpm1*, *avrRpt2*, *avrB*, or *avrRps4* (Varet et al. 2002). *NHL25* expression is also activated upon challenge of Arabidopsis plants that were previously primed by localized Pca infection (Fig. 7A). In Arabidopsis, expression of *NHL25* is only partly induced by salicylic acid (Varet et al. 2002) and it seems that an additional challenging stimulus is necessary to fully activate the gene (Fig. 7A) and counter Pseudomonas infection (Fig. 7H).

## Materials and methods

### Plant growth

Arabidopsis (*A. thaliana*) wild-type (Col-0) and the *nhl25-1* (AT5G36970; SALK_113216) and *npr1-1* (AT1G64280) mutant (both in Col-0 genetic background) were grown on soil in short-day (8 h light, 120 µmol m^−2^ s^−1^) at 20 °C.

### Cultivation of bacteria

Pca and *luxCDABE*-tagged Pca (Pca lux) (Fan et al. 2008) were grown on King’s B medium (20 g L^−1^ tryptone, 10 mL L^−1^ glycerol, 1.5 g L^−1^ K_2_HPO_4_, and 1.5 g L^−1^ MgSO_4_; King et al. 1954) supplemented with 100 µg mL^−1^ streptomycin (Pca) or 25 µg mL^−1^ kanamycin and 100 µg mL ^1^ rifampicin (Pca lux) and 10 g L^−1^ agar. After incubation for 2 d at 28 °C, four or five colonies were transferred to a 250-mL flask containing 50 mL King’s B medium supplemented with 100 mg mL^−1^ streptomycin. The flask was incubated overnight at 28 °C at 220 rpm on a rotary shaker. The next day, the bacterial culture was transferred to a 50-mL plastic tube and centrifuged at 1,800g and 16 °C for 8 min. The supernatant was discarded and the pellet suspended in 50 mL of 10 mM MgCl_2_. After centrifugation at 1,800 g at 16 °C for 8 min, the pellet was suspended in 50 mL of 10mM MgCl_2_. One mL of bacterial suspension was diluted with 10 mM MgCl_2_ to an OD_600_ of 0.0002, resulting in ∼3×10^8^ colony-forming units (cfu) mL^−1^.

### Plant treatment

Using a syringe without a needle, three leaves of 5-week-old Arabidopsis plants were infiltrated with 10 mM MgCl_2_ (mock inoculation) or ∼3×10^8^ cfu mL^−1^ Pca in 10 mM MgCl_2_ (Pca infection). For gene expression analysis, two systemic leaves per plant and treatment were left untreated (no systemic challenge) or challenged by infiltration of tap water at the indicated times after the initial treatment (systemic challenge). Systemic leaves were harvested at 2.5 – 3 h after the systemic challenge and subjected to RT-qPCR analysis as described below.

### SAR assay

Using a syringe without a needle, three leaves of 5-week-old plants were infiltrated with 10 mM MgCl_2_ (mock-inoculation) or ∼3×10^8^ cfu mL^−1^ Pca in 10 mM MgCl_2_. After 72 h, three distal leaves of each plant were infiltrated with Pca lux (∼3×10^8^ cfu mL^−1^) in 10 mM MgCl_2_ (systemic challenge). After another 72 h, leaf discs (0.5 cm diameter) were punched out of inoculated systemic leaves and washed in 10 mM MgCl_2_. Luminescence of Pca lux in leaf discs was measured using CLARIOstar plate reader (BMG LABTECH, Ortenberg, Germany). In the assay, bacterial luminescence correlates with bacterial multiplication (Fan et al. 2008; Gruner et al. 2018).

### Analysis of gene-specific mRNA transcript abundance by RT-qPCR

RNA was isolated from frozen leaves using the TRIZOL method (Chomczynski, 1993). 1 µg of RNA in aqueous solution were subjected to DNase (Thermo Fisher Scientific, Langerwehe, Germany) digestion followed by cDNA synthesis using RevertAid reverse transcriptase (Thermo Fisher Scientific, Langerwehe, Germany). mRNA transcript abundance was determined by RT-qPCR on a C1000 TouchTM Thermal Cycler (CFX 284TM Real-Time System, Bio-Rad, Feldkirchen, Germany) in 384-well Hard-Shell® PCR plates (Bio-Rad, Feldkirchen, Germany) using gene-specific primers (Supplemental Table S2) and iTaq^TM^ SYBR® Green Supermix (Bio-Rad, Feldkirchen, Germany). Data were normalized to the mRNA transcript level of *ACTIN2*.

### STRING database and SUBA5 analysis

The interaction of marker and readout proteins was investigated using STRING database (Szklarczyk et al. 2021) (https://string-db.org). Different levels of stringency were used from high to medium to low confidence (see, the main text). Proteins with medium confidence from the general score were sorted for their gene co-expression score (threshold >0.5) and analyzed for their subcellular localization. A subset of proteins that only interacted with the 14 verified marker and readout genes for priming, was taken and analyzed for their subcellular localization using SUBA5 (Hooper et al. 2022) (http://suba.live/). The SUBA location consensus SUBAcon was used for further subcellular description.

### Statistical analysis

All experiments were repeated at least three times. Statistical significance was determined using GraphPad PRISM (GraphPad Software, San Diego, CA, USA). For experiments with normal distribution Ordinary one-way ANOVA was used, otherwise Kruskal-Wallis test. Changes were considered statistically significant when P>0.05.

## Acknowledgments

We would like to thank Marie Kolvenbach for help with the experiments. Katrin Gruner is thanked for providing Pca lux.

## Author contributions

A.J.S. and U.C. designed the experiments. A.J.S. and F.S. did the experiments. A.J.S and L.F. performed the STRING database and SUBA5 analyses. A.J.S., L.F. and U.C. analyzed the data. U.C. and L.F. wrote the manuscript with input from A.J.S. All authors read and approved the final manuscript.

## Supplemental Material

**Supplemental Figure S1.** SAR to Pca lux is reduced in the *nhl25-2* (GK-340D09) and *npr1* mutant.

**Supplemental Table S1.** ncP, pC, FAIRE, and cP values of priming-readout genes *WRKY6* and *WRKY53*

**Supplemental Table S2.** Gene-specific primer sequences used in RT-qPCR analyses.

**Supplemental Dataset S1.** Co-expression of genes.

## Funding

### Conflict of interest statement

We declare we have no conflict of interest. Uwe Conrath is a cofounder of AgPrime GmbH.

## Data availability

The authors confirm that all experimental data are available and accessible via the main text and/or the Supplemental Material.

